# Assessing the reliability of species distribution projections in climate change research

**DOI:** 10.1101/2020.06.10.143917

**Authors:** Luca Santini, Ana Benítez-López, Luigi Maiorano, Mirza Čengić, Mark A.J. Huijbregts

## Abstract

**Aim:** Forecasting changes in species distribution under future scenarios is one of the most prolific areas of application for species distribution models (SDMs). However, no consensus yet exists on the reliability of such models for drawing conclusions on species distribution response to changing climate. In this study we provide an overview of common modelling practices in the field and assess model predictions reliability using a virtual species approach.

**Location:** Global

**Methods:** We first provide an overview of common modelling practices in the field by reviewing the papers published in the last 5 years. Then, we use a virtual species approach and three commonly applied SDM algorithms (GLM, MaxEnt and Random Forest) to assess the estimated (cross-validated) and actual predictive performance of models parameterized with different modelling settings and violations of modelling assumptions.

**Results:** Our literature review shows that most papers that model species distribution under climate change rely on single models (65%) and small samples (< 50 presence points, 62%), use presence-only data (85%), and binarize models’ output to estimate range shift, contraction or expansion (74%). Our virtual species approach reveals that the estimated predictive performance tends to be over-optimistic compared to the real predictive performance. Further, the binarization of predicted probabilities of presence reduces models’ predictive ability considerably. Sample size is one of the main predictors of real accuracy, but has little influence on estimated accuracy. Finally, the inclusion of irrelevant predictors and the violation of modelling assumptions increases estimated accuracy but decreases real accuracy of model projections, leading to biased estimates of range contraction and expansion.

**Main conclusions:** Our study calls for extreme caution in the application and interpretation of SDMs in the context of biodiversity conservation and climate change research, especially when modelling a large number of species where species-specific model settings become impracticable.

## 1. Introduction

Understanding how climate shapes species distribution and how range shifts may be driven by future climatic change is more urgent than ever. In the last thirty years, studies aimed at developing, improving and applying species distribution models (SDMs) have proliferated (Araújo et al. 2019), and forecasting changes in species distribution under future scenarios is one of the most popular areas of application for SDMs today (Thuiller et al. 2011, Schloss et al. 2012, Newbold 2018). In SDM-based climate change forecasting studies, models are trained on current data and used to predict the probability of presence under present and future conditions. Models’ predictions are often binarized to assess whether a species distribution is expected to shift, contract or expand (Newbold 2018). Although many modelling techniques require presence and absence data, many models are fitted using presence-only data, i.e., contrasting presences with random pseudo-absences, or background points, that represent available conditions (Guillera-Arroita et al. 2015). The predictive performance of these models is commonly assessed by randomly splitting the dataset into training and testing, and fitting the model on the training dataset and validating it on the testing dataset using discrimination metrics such as the True Skill Statistic (TSS) or the Area Under the Curve (AUC). While several authors have warned about the challenges and uncertainties of projecting future species distribution (Dormann 2007, Peterson et al. 2018), only few studies have tested model performance with empirical data, reporting mixed results (Araujo et al. 2005, Rapacciuolo et al. 2012, Morán-Ordóñez et al. 2017, Sofaer et al. 2018).

The literature on SDMs has grown very quickly and extensively, with papers adhering to different schools of thoughts and supporting the use of one or another technique (see Norberg et al. 2019 for an overview), suggesting different validation measures (e.g. Allouche et al. 2006, Leroy et al. 2018) or approaches (e.g. testing on spatially independent data; Bahn and McGill 2013). Additionally, a number of studies made different conclusions about the minimum number of presence points needed (Stockwell and Peterson 2002, Hernandez et al. 2006, Wisz et al. 2008, van Proosdij et al. 2016), area of sampling of background points (e.g. VanDerWal et al. 2009, Anderson and Raza 2010, Elith et al. 2010, Barve et al. 2011), or choice of environmental predictors and approaches to reduce collinearity (see Fourcade et al. 2018 for an overview). This can make it challenging and disorientating for people that approach the field of SDM for the first time. The existence of modeling software that make the application of these models easier and more accessible to people with limited modelling background (e.g. MaxEnt Phillips et al. 2006), may be counterproductive, as running an SDM in one of these software may appear simpler than it is. This is particularly worrying considering that SDMs are largely used to inform conservation science (Newbold 2018, Manish and Pandit 2019).

Recently, a number of experts have delineated a set of best practices, and shown that many studies still apply inconsistent approaches that do not adhere to the best standards (Araújo et al. 2019). This generates a self-perpetuating problem, because published papers create a precedent, and are used to justify modelling choices in new papers. For example, while several authors argued that models’ predictors should be chosen considering the biology of the species (Araújo and Guisan 2006, Austin and Van Niel 2011), it has become a common practice to include all bioclimatic variables excluding collinear variables using automatic procedures irrespective of species-specific biological considerations (e.g. Manish and Pandit 2019), increasing the risk of detecting spurious relationships (Synes and Osborne 2011, Fourcade et al. 2018). Worryingly, it has been shown that non-biologically relevant predictors can contribute to increase the predictive ability of the models (Fourcade et al. 2018), so discrimination accuracy metrics may suggest a very good model while the relationships estimated do not have a biological meaning (Journé et al. 2019, Warren et al. 2020). Spurious relationships become particularly problematic when the model is projected to new areas or environmental scenarios (Heikkinen et al. 2012, Bahn and McGill 2013, Yackulic et al. 2013, Merow et al. 2014). Similarly, methodological papers that suggest less demanding requirements can become preferred and widely cited references, reinforcing the trend. For example, van Proosdij et al. (2016) concluded that 14 to 25 observations may be sufficient to run species distribution models. This is now often cited to justify the use of small sample sizes (e.g. Carlson et al. 2017, Chen et al. 2017) despite previous recommendations suggesting a minimum of 50 points (Stockwell and Peterson 2002, Hernandez et al. 2006, Wisz et al. 2008).

The extent to which SDMs perform adequately also depends on the degree to which modelling assumptions are met. SDMs are often fitted on opportunistically collected data that violate the assumption of random sampling (e.g. Guillera-Arroita et al. 2015). Furthermore, the present distribution of species is rarely in equilibrium with the environment, meaning that species only occupy a portion of the fundamental niche, not only because of biotic (e.g. competition or predation) or dispersal constraints (e.g. physical barriers, limited dispersal abilities; Soberon and Peterson 2005), but also because they may have recently contracted their distribution due to human influence (e.g. Varela et al. 2009, Di Marco and Santini 2015, Faurby and Svenning 2015) or stochastic events. This problem has often been discussed in the literature in relation to the inferences made (Varela et al. 2009, Maiorano et al. 2013, Martínez-Freiría et al. 2016, Faurby and Araújo 2018). Yet, methodological papers aimed at assessing optimal settings to run species distribution models typically assume ideal conditions (e.g. van Proosdij et al. 2016).

Models used for future projections to inform conservation need to adhere to even higher standards than those used for present predictions (Sequeira et al. 2018). In fact, while a model used for predicting current distribution can still provide meaningful predictions even though the inferred relationships are wrong (Fourcade et al. 2018, Warren et al. 2020), relationships need to be realistic in order to make meaningful predictions to different conditions. However, although guidelines for transferability have been provided (Sequeira et al. 2018), it is common to validate models on present data and assume they perform equally well for future predictions. While a number of studies have discussed and tested the influence of multiple sources of uncertainty on the predictive accuracy of SDM predictions under present conditions (e.g. Wenger and Olden 2012, Vale et al. 2014, Fourcade et al. 2018, Fernandes et al. 2019), to our knowledge, no study has yet tested the reliability of both present and future predictions while considering the effects of different modelling settings and several violations in model assumptions simultaneously.

In this study we first provide an overview of common practices in the field by reviewing the papers published in the last 5 years. We focused on the sample size used, choice and selection of environmental predictors, types of models employed, the sampling approach of background (or pseudo-absence) points, and the method used for binarization of model outputs. Then, we employed a virtual species approach (Zurell et al. 2010, Meynard et al. 2019) to assess the contribution of different modelling settings and violation of assumptions to the predictive accuracy and projected responses to climate change of SDMs for three commonly applied model algorithms (GLM, MaxEnt and RandomForest). Our approach allows validating model predictions against the virtual “reality”, therefore estimating true model predictive accuracy. We generated 50 virtual species distributions, fitted SDMs under different conditions, and assessed the discrimination ability of present and future model predictions against the real distribution. We also compared this predictive ability with that estimated using a cross-validation (split-plot) approach, which is the most common way of assessing model discrimination accuracy in most SDM studies. We systematically assessed the combined effect of 1) the number of presence points (i.e. sample size), 2) the geographic extent over which background points are drawn, 3) the number of biologically relevant (i.e. true niche axes) and irrelevant predictors (i.e. spurious correlates), 4) the species prevalence (proportion of study area occupied by the species), 5) the sample prevalence (proportion of presences over background points), 6) the proportion of niche filling (the degree to which the species is at equilibrium with the environment), 7) and the spatial bias in presence points. We then assessed model predictions using two common discrimination metrics: the AUC and the TSS.

## 2 Methods

### 2.1 Literature review

We conducted a literature review on common practices in SDM papers that projected models to a different time period (past or future). We queried Web of Science and focused on papers published in the last 5 years (2015-2019) to reflect the most recent trends in the field. We randomly selected 50 papers per year for a total of 250 papers. From each paper we extracted the following information: sample size, occurrence data type (e.g. presence-only vs. presence-absence), models used, the variable selection criteria, and whether probabilistic output were binarized or not. A detailed description of the literature search and data extraction is presented in Supplementary material Appendix 1.

### 2.2 Environmental variables

We obtained 19 bioclimatic variables from CHELSA (http://chelsa-climate.org; Karger et al. 2017) at 0.1 degree resolution (∼11 km) for the present and for the future (year 2050) RCP 8.5 taking the median over all the General Circulation Models (GCM). We also downloaded the human footprint index for 2009 from https://wcshumanfootprint.org/ (Venter et al. 2016).

### 2.3 Virtual species

We generated 50 virtual species using the ‘virtualspecies’ R package (Leroy et al. 2016). For each virtual species, we first determined the study area by generating a random extent between 3 and 10 decimal degrees (∼330-1100 km) in both longitude and latitude centered around a random location in the globe (Fig. 1). We then selected 6 random bioclimatic variables and sampled their values within the extent using 100 random points. We used the mean and standard deviation estimated for the 6 bioclimatic variables to generate the niche tolerance for the virtual species.

**Fig. 1.**
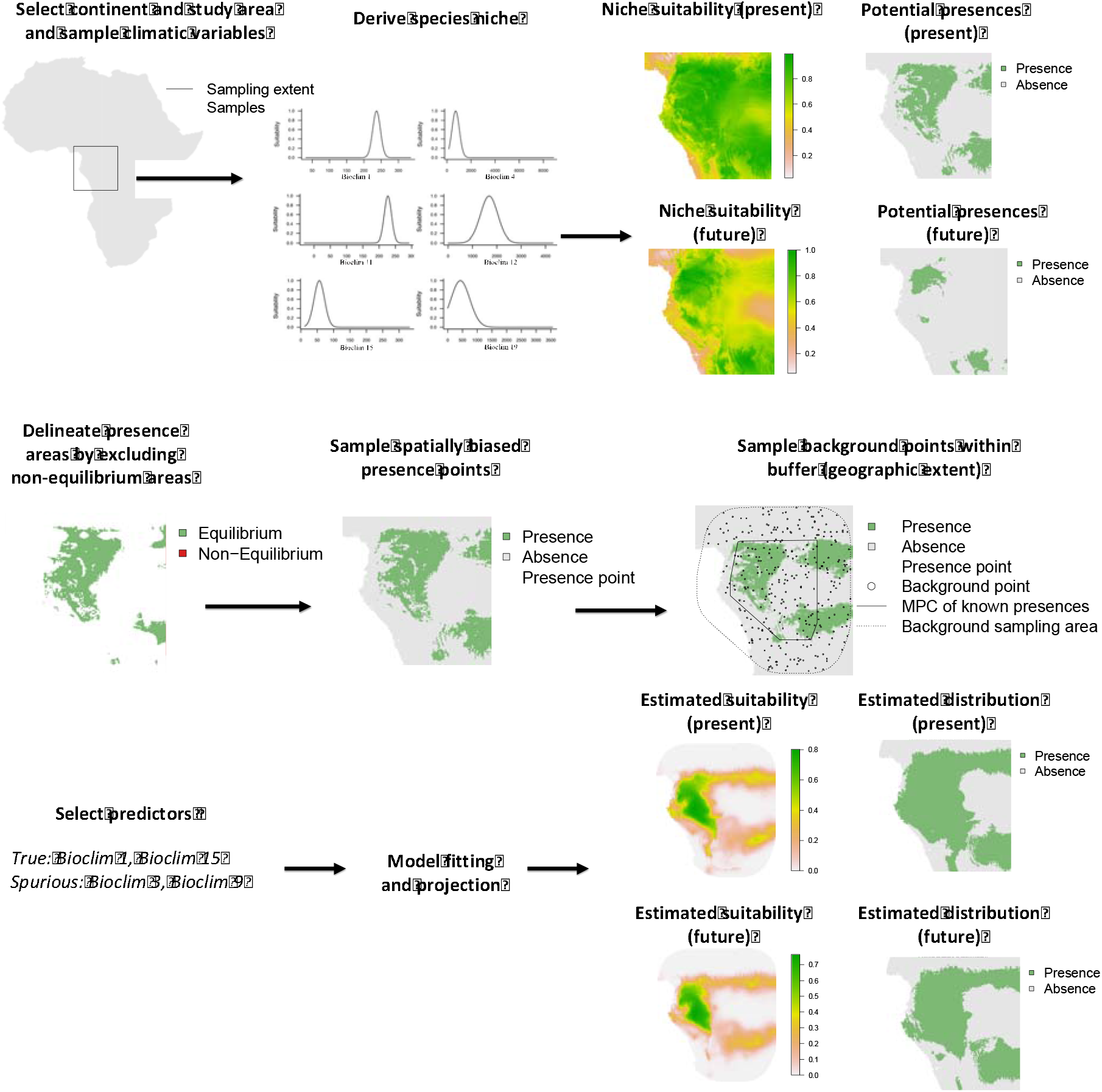
Modelling steps taken to generate virtual species and fit and project the species distribution model.

We then projected the niche within the study area for present and future conditions and defined the occupied area using a threshold sampled randomly between the 0.2 to 0.8 quantiles of the suitability values in the study area. The threshold is meant to represent the values above which the species can survive and is assumed to be present for the validation of the distribution models (see section 2.5). Note that the virtual species can potentially be present outside this study area in environmentally analogous conditions, but we assume that the species is either limited by dispersal, absent because of biotic interactions, or its presence outside the study area is simply unknown to the modeller.

### 2.4 Scenario settings

For each virtual species, we fitted species distribution models using different cross-combinations of the settings presented in Table 1. To assess the influence of sample size, we sampled random presence points (10, 25, 50, 100, 250, 500 and 1000 points) within the distribution area of the species. Presences were sampled randomly and not as a function of niche suitability values as there is no evidence that species abundance increases with niche suitability (Dallas and Hastings 2018) and observation probability is often also a function of other factors such as vegetation structure or human presence. Then, to assess the influence of the geographic extent, we fitted a minimum convex polygon (MCP) around presence areas to generated a buffer expressed as percentage increase of the MCP, which delimited the geographic extent within which the background points were sampled (0%, 100%, 500%, 5,000%, 50,000%; the latter often resulting in the entire continent). We set the number of background points depending on the number of presence points and the level of sample prevalence. We used three sample prevalence: 0.01, 0.1 and 1. Not all background points, however, could always be sampled depending on the selected geographic extent (i.e. insufficient number of cells), leading to variable sample prevalence values.

**Table 1.** Summary of treatments considered for fitting the species distribution models on virtual species.

| Treatment | Values |
| --- | --- |
| Presences | 10, 25, 50, 100, 250, 500, 1000 |
| Sample Prevalence | 0.01, 0.1, 1 |
| Buffer (%) | 0, 100, 500, 5000, 50000 |
| Bias (%) | 33, 66, 100 |
| Niche filling (%) | 33, 66, 100 |
| Relevant predictors | 0, 3, 6 |
| Irrelevant predictors | 0, 3, 6 |

In each model, we used a total number of predictor variables between 3 and 12. To assess the influence of biologically relevant and irrelevant predictors of species presence, we sampled none, 3, or 6 relevant bioclimatic predictors (those describing the true species niche), and none, 3 or 6 irrelevant bioclimatic predictors (not describing the niche) from the other 13 bioclimatic variables (Table 1). Combinations yielding 0 predictor variables were not considered. We tested collinearity using a stepwise VIF selection for the environmental variables in the training dataset and only retained variables with VIF<3 (Zuur et al. 2010), so the final number of biologically relevant or irrelevant predictors could be different from multiples of 3. As a measure of model transferability, we estimated the Multivariate Environmental Similarity Surface (MESS; Elith et al. 2010) between the present and future set of environmental variables used in the model fitting.

We also considered violations of two important assumptions underlying SDMs that are common in real study cases: non-equilibrium with the environment (niche filling) and non-random sampling of presence points that results in a bias along an environmental gradient. Decreasing proportions of niche filling were simulated by only sampling presence points below a given quantile (0.33, 0.66, 1) of the human footprint index values within the study area (Table 1). This mimics a scenario where a species is potentially present (given climatic conditions) and yet absent because of human impact. Note that species may be in disequilibrium with the environment for different reasons (e.g. biotic interactions, dispersal limitations) but the result would be similar. For simplicity we restrict our analyses to the case where species are not an equilibrium because of human impact.

Environmental bias was simulated by randomly sampling one of the biologically relevant bioclimatic predictors used in the distribution model and sampling presences only below a given quantile (0.33, 0.66, 1) of the distribution of environmental values (Table 1). This represents the situation where the species has only been observed under certain conditions (i.e. sampling bias correlates with environmental gradients), therefore potentially biasing the estimation of the species niche. When no biologically relevant variable was included, an irrelevant predictor was selected instead.

The full set of combinations of settings in Table 1 corresponded to 7560 models; to reduce the computational effort we sampled 500 model settings from the multidimensional space using a conditional Latin hypercube approach (Minasny and McBratney 2006), which ensured that the subset of models is representative of the real variability occurring in the original 7560 models.

### 2.5 Model fitting and validation

We used this synthetic dataset to fit three distribution model algorithms: MaxEnt (using a ‘cloglog’ transformation and linear and quadratic feature classes), Generalized Linear Model (GLM, with a stepwise model selection based on AIC including linear and quadratic terms and weights set for equal sample prevalence), and Random Forest (with stratified sampling, 500 trees, and an ‘mtry’ parameter equal to the rounded square root of the number of predictor variables). For each, we run a repeated split sample cross-validation by splitting the dataset into training (80%) and testing datasets (20%) 10 times. We estimated model discrimination accuracy with the Area Under the Curve (AUC) and the True Skill Statistic (TSS) (Lawson et al. 2014). Then, we fitted the model using the full sample, and binarized the predictions into presence-absence by using the threshold that maximized TSS. We estimated contraction and expansion areas by overlaying the binary predictions for the present and the future. Finally, we validated the model predictions for the present, future, and areas of contraction and expansion against the virtual reality using the same discrimination metrics. This validation was performed within the area of background point sampling. We matched the predicted probabilities with the true presences and absences of the virtual species to estimate the true AUC, and the predicted presences and absences from the binarized model with the true presences and absences of the virtual species to estimate the true TSS using the threshold that maximized TSS on the testing dataset. By doing this, we were able to both 1) estimate model discrimination accuracy mimicking a typical ecological modeller, and 2) quantify the real model discrimination accuracy by comparing the model to the virtual reality.

### 2.6 Evaluation of model settings

As a post-processing step, we used a Random Forest regression to estimate the influence of different modelling settings and confounding factors on the discrimination accuracy of the three distribution model algorithms. We fitted a Random Forest with 1,000 trees to each species using all discrimination performance metrics (TSS and AUC, both estimated and true for the present and the future) and estimated changes in distribution (% of range contraction and expansion) as dependent variable (one model per dependent variable), and the values of each treatment (number of presences, sample prevalence, species prevalence, environmental similarity, number of relevant predictors, number of irrelevant predictors; % buffer, degree of bias in sampling points, niche filling proportion) as independent variables. The ‘mtry’ parameter in the random forest model was set to the number of predictors divided by 3 (Breiman 2001).

We then estimated the relative importance by permutation and partial response curves for each predictor per species, and then averaged all relative importance estimates and partial response curves per variable across all species. The estimated relative importance values were transformed to percentages (rescaled to 100) for interpretability. Confidence intervals for both variable importance and partial response curves were estimated from the standard error of the mean across species models.

### 2.7 R packages

All analyses were computed in R v. 3.5.3 (R Core Team 2018) using the packages ‘virtualspecies’ (Leroy et al. 2016), ‘raster’ (Hijmans and van Etten 2014), ‘PresenceAbsence’ (Freeman and Moisen 2015), ‘dismo’ (Hijmans et al. 2017), ‘rgeos’ (Bivand and Rundel 2013), ‘pROC’ (Robin et al. 2013), ‘usdm’ (Naimi 2013) and ‘GISTools’ (Brunsdon and Chen 2014) for generating virtual species and fitting species distribution models, and ‘clhs’ for the conditional Latin hypercube sampling (Roudier 2011). We used the R package ‘randomForest’ (Liaw and Wiener 2002) and ‘ranger’ for fitting Random Forest models (Wright and Ziegler 2017), ‘maxnet’ package to fit MaxEnt (Phillips 2017), and ‘pdp’ for estimating the partial response curves (Greenwell, Brandon 2019). The codes used for the analyses of this paper are available as part of the supplementary materials.

## 3. Results

### 3.1 Common practices in SDMs

Among 250 papers reviewed, 92 included correlative species distribution models projected to different times, and therefore were deemed relevant for our scopes (Table S1). Based on our sample and using a bootstrapping approach, we estimated that the total number of papers published between 2015 and 2019 that matched this criterion is 1194-1665 (95CI), indicating that we sampled approximately between 5.5 and 7.7% of the total (Appendix 1).

Most of the papers inspected included models fitted on relatively small sample sizes (N < 50; Fig. 2a), with only 18.4% including minimum samples larger than 50 and 16.1% not reporting the sample size used. More than 50% of the papers included all bioclimatic variables with no biological justification (Fig. 2b). Among these papers, in ∼50% of the cases the authors reduced the number of variables using automatized approaches based on correlations or best fit to the data. A smaller number of studies selected variables a priori, some of which did not provide a justification for this choice (7.8%). Most of the studies used a single model (Fig. 2c,d), with MaxEnt being the most common algorithm used (78.3%), followed by linear models (GLM = 30.4%; GAM = 26.1%) and machine learning models (RF = 27.2%; GBM = 20.7%) (Fig. 2c). The majority of studies did not include real absences but used pseudo-absences, background data, or presence-only methods (i.e. climatic envelopes; 84.8%; Fig. 2e). A large proportion of papers using pseudo-absences or background points did not report the area of sampling (48.7%), while others used a variety of different approaches, the most common being sampling randomly across the pre-defined study area (Fig. 2f). Finally, most studies binarized the continuous probability outputs based on discrimination metrics (e.g. max TSS or equal sensitivity and specificity; 73.9%), almost one quarter of the studies reported the continuous output (22.8%), and a small percent (3.3%) categorized probabilities into multiple arbitrary categories (e.g. 0.3, 0.6 and > 0.6; Fig. 2g).

**Fig. 2.**
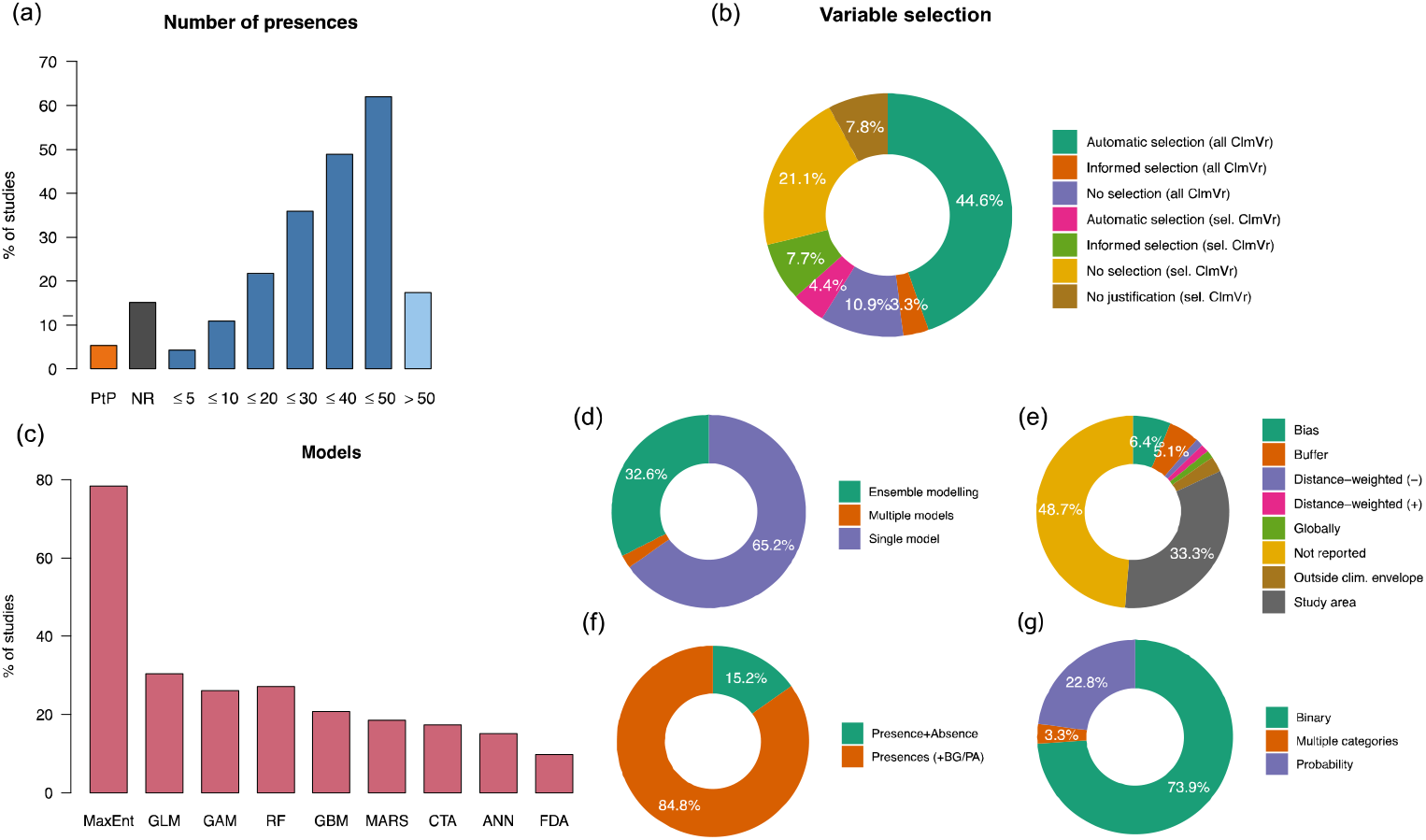
Summary of the literature review. (a) Number of presences used in the models (minimum among species if multiple species were modelled); PtP = Polygons converted to presence points; NR = sample size not reported; (b) Variable selection approach. all ClmVr = all climatic variables were considered; sel. ClmVr = a subset of climatic variable was considered; Automatic selection = collinear variables were excluded using automatized approaches based on correlations, variance inflation factors, or best fit to the data; Informed selection = collinear variables were excluded based on expert opinion; No selection = Collinear variables were not excluded, or no collinearity was not found or reported; No justification = no justification provided for the rationale underlying the subset of variables chosen; (c) Percentages of studies using MaxEnt, generalized linear models (GLM), generalized additive models (GAM), random forests (RF), generalized boosted trees (GBM), multivariate adaptive regression splines (MARS), classification trees (CTA), artificial neural networks (ANN) and flexible discriminant analysis (FDA). Other models used in a minority of instances are not reported here (see Table S1); (d) Percentage of studies using one or multiple models, or ensemble modelling approach; (e) Pseudo-absences or background points sampling approach; Bias = sampling that mimics sampling bias; Buffer = random sampling within a buffer around presence points; Distance-weighted = Sampling with higher intensity near (-) or far from (+) presence points; Globally = Random sampling globally; Outside climate envelope = Beyond climatic conditions observed for presence points; Study area = random sampling within a pre-defined study area; (f) Percentage of studies using presences only, presences + background points / pseudo-absences, and those using presences and real absences; (g) Percentage of studies binarizing the probabilities into suitable/unsuitable, or in multiple arbitrary categories, or not applying any form of binarization.

### 3.2 Reliability of climate change predictions

The three algorithms showed a consistent pattern across the two scenarios and accuracy metrics. The predictive accuracy of models’ predictions estimated by cross-validation was consistently above the typically accepted performance thresholds (AUC=0.7, TSS=0.5; Landis and Koch 1977, Swets 1988), and higher than the true predictive accuracy for present, future predictions, and contraction and expansion areas (Fig. 3). However, the accuracy of binary predictions (TSS) was substantially lower than that measured for continuous predictive outputs (AUC), suggesting that the binarization of relative probabilities of presence decreases models’ predictive ability considerably (Fig. 3). Models’ predictions for the future and contraction and expansion areas showed lower predictive performance (Fig. 3a), especially when binarized (Fig. 3b).

**Fig. 3.**
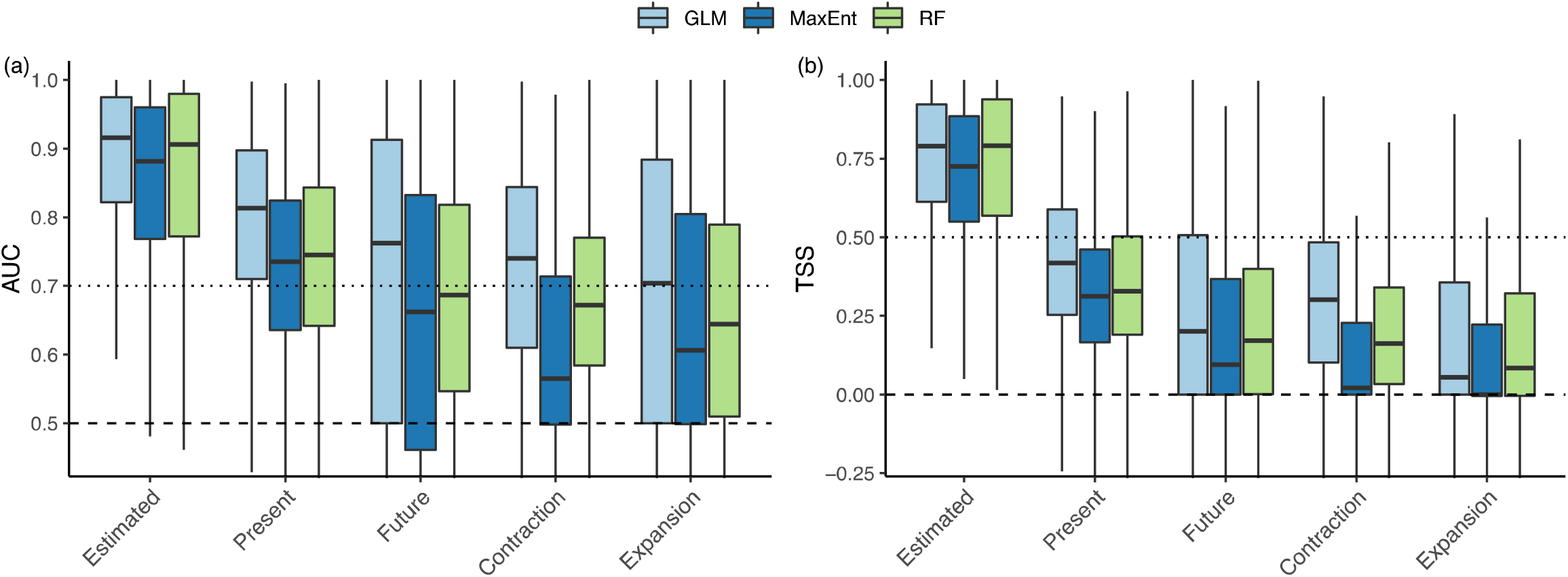
AUC (a) and TSS (b) of the models fitted. Estimated = Estimated through internal cross-validation; Present and Future = True value validated against virtual reality for present and future; Contraction and Expansion = True value validated against virtual reality for predicted contraction and expansion areas; Dashed line = null expectation (no better than random); Dotted line = Value typically considered as “good” performance thresholds. The box edges are the 25th and 75th percentiles of the distribution, and whiskers 1.5 the inter-quartile range.

Under optimal modelling settings (e.g. large sample size, relevant predictors, no violation of assumptions regarding niche filling and unbiased sampling), models performed relatively well according to AUC (Fig. S1a), but poorly when considering binary outputs (Fig. S1c). On the contrary, under poor modelling settings and conditions (small sample size, irrelevant predictors, violation of the main assumptions), the estimated predictive abilities remained high, but the true predictive abilities dropped considerably, especially when predictions were binarized into presence-absence (Fig. S1b,d). TSS and AUC were highly correlated, but while high TSS always corresponded to high AUC, the opposite was not always true (Fig. S2).

### 3.3 Determinants of estimated predictive ability

The importance and effect of different factors on the estimated predictive accuracy by cross-validation was qualitatively similar when using TSS or AUC (Fig. S2-S8). The most important predictors of estimated predictive accuracy were species prevalence, the environmental gradient sampled (inverse of environmental bias), and the geographic extent of background point sampling (Fig. 4). Additionally, sample prevalence was important for Random Forest, and the number of presence points for GLM (Fig. 4). Predictive accuracy decreased with increasing species prevalence and decreasing environmental gradient sampled (i.e. increased with environmental bias), and increased with increasing geographic extent sampled (Fig. S3-S8). The number of presence points had a positive effect when fitting GLM and MaxEnt models, but had little effect when using Random Forests. Sample prevalence a had positive effect in Random Forests, and weakly negative in the other two models. The number of relevant and irrelevant predictors had a weak but positive effect regardless of the model (Fig. S3-S8).

**Fig. 4.**
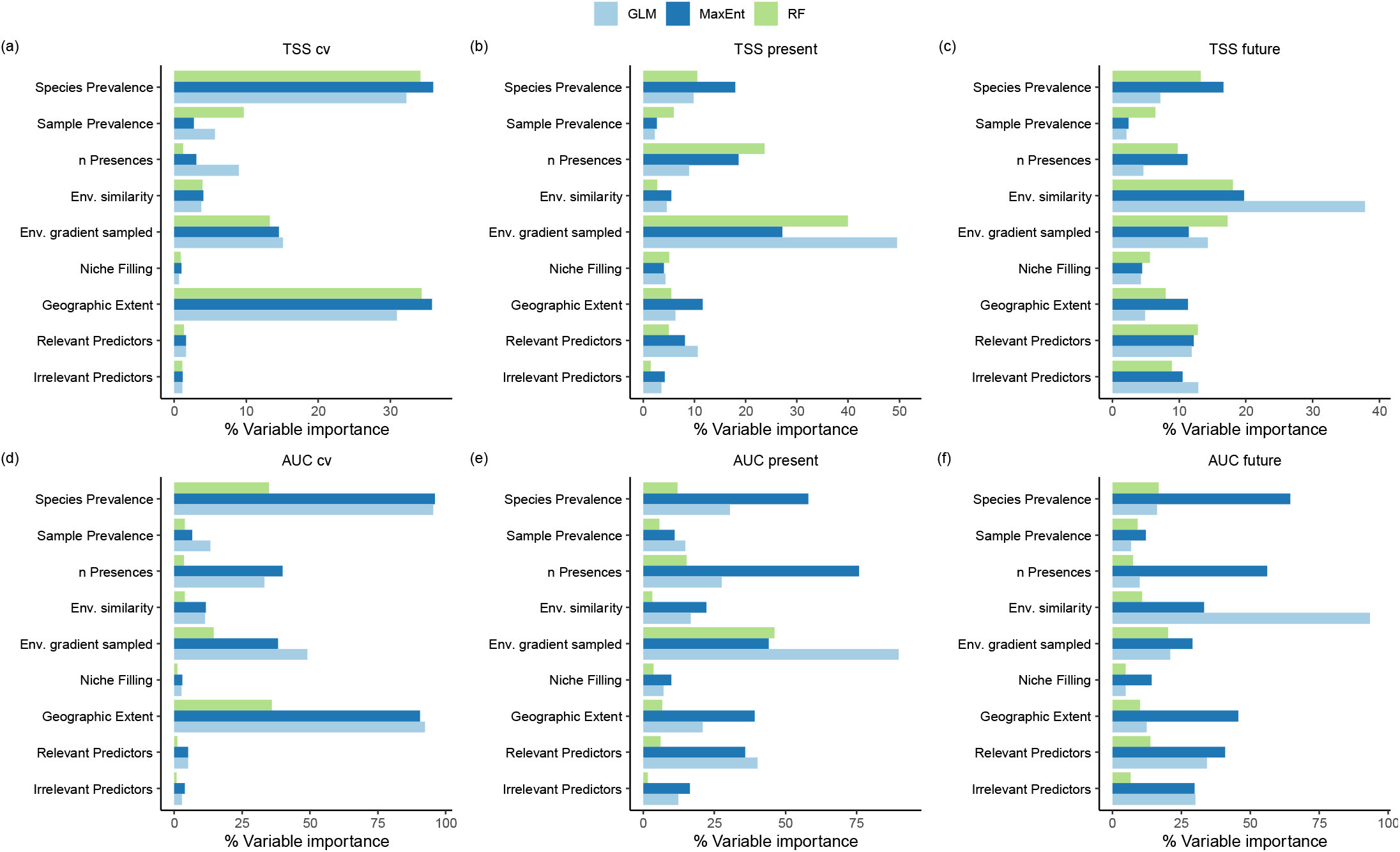
Relative variable importance of different settings and conditions on the TSS (a, b, c) and AUC (d, e, f) estimated by cross-validation (a, d), and measured against virtual reality for the present (b, e) and future predictions (c, f). Relative importance values are rescaled to 100 for each species. Bars represent the mean over all virtual species and error bars the standard error around the mean.

### 3.4 Determinants of true predictive ability

The true predictive accuracy (i.e. measured against the virtual reality) of the models for the present was mostly affected by species prevalence, the number of presences, and the environmental gradient sampled. The geographic extent was also important when fitting MaxEnt models (Fig. 4). Both the number of presence points and the environmental gradient sampled had a positive influence on predictive accuracy, geographic extent had weak positive effect, and the species prevalence a negative effect (Fig. S3-S8).

The number of biologically relevant and irrelevant predictors, and niche filling, were relevant for present predictions, but became especially influential for the predictive accuracy of models projected into the future, with the number of relevant predictors and niche filling increasing predictive performance, and the number of irrelevant predictors decreasing predictive performance (Fig. S3-S8). An important predictor of the predictive accuracy of future projections was the degree of environmental similarity between the present and future environmental conditions of the study area (Fig. 4, Fig. S3-S8).

Species with high prevalence were more likely to expand and less likely to contract the range. However, a number of additional factors contributed to these estimates (Fig. S9-S12), such as the number biologically relevant and irrelevant predictors, showing a positive effect on contraction and expansion estimates in GLM and MaxEnt, and a negative effect on contraction areas in random forest (Fig. S10-S12). The environmental similarity between present and future conditions yielded a negative effect on contraction and expansion areas, but showed non-linearity for contraction areas estimated by GLM and Random Forest models. Violation of equilibrium and random sampling assumption also contributed to increase range contraction and expansion estimates (Fig. S10-S12).

## 4. Discussion

In this paper we report on common practices in SDM and use this information to assess the effects of these practices on the predictive accuracy of SDMs, and thus, on the reliability of future climate-induced range shifts. Our literature review points out that a large part of papers that model species distribution under climate change rely on single models (typically MaxEnt), include models fitted on very small samples, use presence-only data, and typically binarize models’ output to measure range shift, contraction or expansion. Consistently with previous analyses (Araújo et al. 2019), it also highlighted how poor modelling practices are common in the literature, especially in relation to the use of very small samples, lack of ecological considerations in the selection of model predictors, and non-reporting of fundamental information on background sample selection and study area (Zurell et al. 2020). When exploring the influence of these practices on the predictive accuracy using a virtual species approach, we found out that the estimated discrimination capacity by TSS and AUC does not reflect the actual predictive ability of SDMs, and tends to be over-optimistic compared to the real model performance when predicted under present conditions, and especially when projected to future (different) conditions. The ability of models to discriminate presences from absences as measured by the TSS is particularly low, even under optimal model settings, good ecological knowledge of the species climatic requirements, and modelling assumptions are fully met. The extent to which predictions are reliable depends on a number of model parameters (e.g. number of presence points), actual proportion of species distribution within the geographic extent (species prevalence), our degree of knowledge of the species ecology (predictor variables included in the model), and difference between present and future environmental conditions. Under optimal settings and a good ecological knowledge of the species climatic requirements, future predictions show low discrimination ability, whereas under non-optimal settings, predictions may not be better than random. Ultimately, our results suggest that irrespective of the estimated performance, we may be unable to make meaningful future predictions for many species, and even when we can, binarization of models’ outputs should be avoided. Based on our results, we elaborate in the following paragraphs on guidelines and recommendations for good modelling practices when fitting SDMs.

### 4.1 Aim for large sample sizes

An important determinant of predictive accuracy that is often undervalued is sample size. Previous studies suggested a minimum of 50 points (Stockwell and Peterson 2002, Hernandez et al. 2006, Wisz et al. 2008), and van Proosdij et al. (2016) suggested even fewer were needed.

However, these studies assessed the number of points needed under optimal conditions where the modeller uses biologically relevant environmental predictors, points are sampled randomly, and species are in equilibrium with the environment; or used real species (therefore estimating accuracy with testing data, e.g. Wisz et al. 2008). Our results show that while sample size has a little influence on the estimated (cross-validated) accuracy, it is one of the most important predictors of true accuracy. The relationship with sample size is asymptotic, and tends to stabilize around 200-500 points. We must stress, however, that no magic number exists, and these values are contingent on the settings in our simulation (e.g. the number of predictor variables used in the models). Because we are rarely aware if the predictor variables are directly linked to species ecology, or if the species is in equilibrium with the environment or presence points are biased, one should always aim for the largest possible sample. This may be impracticable for many species, that are either poorly known, or narrow ranged. In the absence of biological information on e.g. species’ thermal tolerance, it is hard to say, however, if species that are narrow ranged are specialist of specific climate conditions, or are in disequilibrium with the environment. This second case likely would result in an under-estimation of niche tolerance and over-prediction of range contraction under climate change (Araújo and Pearson 2005, Martínez-Freiría et al. 2016, Faurby and Araújo 2018). In these cases, alternative conservation assessments should be considered when possible. Projecting SDMs trained on insufficient samples does not improve our knowledge in any meaningful way and may actually be detrimental.

### 4.2 Behold sample prevalence, not the absolute number of background points

Many SDM studies using presence-only data sample a large number of background points or pseudo-absences (e.g. 10,000), often citing Barbet-Massin et al. (2012) or Phillips and Dudík (2008) as supporting reference. However, Barbet-Massin et al. did not test MaxEnt, and showed important differences between algorithms. In turn, Phillips and Dudík (2008) tested MaxEnt but they report their results for many species with different numbers of presence points. Hence, the positive relationship between AUC and the number of background points they found should be interpreted carefully as it is mediated by sample prevalence. A recent study concluded that the number of background points depends on the modelling technique used (Liu et al. 2019), with accuracy in MaxEnt stabilizing above a few hundreds of background points, and large numbers being only relevant for common species with small samples of training presences. Our results show that GLM and MaxEnt work best when sample prevalence is very low, supporting the practice of sampling a large number of background points or pseudo-absences compared to the number of presences.

However, matching the findings by Barbet-Massin et al. (2012), we found that Random Forest models perform best with high sample prevalence. This reinforces the notion that no rule of thumb exists and settings should be model- and sample-specific, which is often ignored in ensemble forecasting approaches that fit all models on the same dataset (e.g. Avalos and Hernández 2015, Sales et al. 2017).

### 4.3 Choose predictors carefully

The number and quality of predictors does not seem to have a clear effect on estimated accuracy, if any, increasing the number irrespective of the true underlying relationship, tends to deceitfully increase estimated performance, and increase or decrease the estimated range contraction and expansion. Choosing biologically meaningful predictors may not be particularly problematic when predicting to present conditions (Fourcade et al. 2018), but it becomes a serious issue when the model is transferred in space or time (Wenger and Olden 2012, Sequeira et al. 2018). Here we considered an optimistic scenario where only 6 climatic variables influence species distributions. In reality, there might be many biologically relevant variables that determine or influence the distribution of a species, but our results suggest that when model is projected under different conditions is better to aim for few variables for which we have clear biological expectations than many variables with unclear effects on the species’ distribution (Araújo and Guisan 2006, Austin and Van Niel 2011).

### 4.4 Geographic extents should accommodate the purpose of the study

Previous studies suggest sampling background points in areas that are potentially accessible to the species (e.g. biome or continent) (Araújo et al. 2019) or considering the historical biogeography of the species (Barve et al. 2011, Merow et al. 2013, Cooper and Soberón 2018). This is meant to allow a fair comparison between what is used and what is available. Sampling over large areas tend to inflate estimated predictive accuracy, whereas the effect on true predictive accuracy of present and future predictions is inconsistent across metrics (positive for AUC and negative or flat for TSS) and models. This suggests that the most appropriate geographic area for sampling background points varies across species and it should be tailored to the objective of the study. Setting a biologically meaningful sampling area requires a deep knowledge of species ecology (e.g. dispersal distance, physical and biotic barriers) and biogeography (e.g. historical distribution), which is unavailable for most species, an important future avenue of research may be delineating rules of thumbs that tend to improve accuracy.

### 4.5 Noise is inevitable

An important driver of the variation in model performance is species prevalence (Leroy et al. 2018). Our results concur with previous studies showing that generalist species are harder to predict than specialist species (Evangelista et al. 2008). However, “generalist” and “specialist” are relative terms in the context of species distribution models, as they are defined based on the geographic extent being sampled. Species prevalence over the geographic extent is something we are unaware of in real study cases, and will always be an unknown factor that affects our predictive ability (Leroy et al. 2018). In this sense, we should aim to optimize other model settings that can be controlled for, such as the choice of predictors, the sample prevalence or having a biologically plausible geographic extent.

Our results also show that when future environmental conditions are very dissimilar from present conditions, model’s projection tend to perform poorly. While entirely expected as model predictions extrapolate beyond the model domain (Elith et al. 2010), this is in a way paradoxical. In fact, the more dissimilar future conditions will be, the more species are expected to shift their distribution range and projections becomes important to inform conservation science. Our results not only corroborate previous studies emphasizing the importance of identifying extrapolation areas for highlighting projection uncertainty (Elith et al. 2010, Owens et al. 2013), but also indicate that forecasting accuracy decreases substantially an already low predictive performance.

### 4.6 Violation of modelling assumptions provides a false sense of accuracy

Species distribution models rely on the assumptions of random sampling and species equilibrium with the environment. Worryingly, our results show that when these two assumptions are not met, the estimated accuracy by cross-validation can be inflated, therefore giving the false impression that the model performs well. The extensive use of citizen science data in SDMs make models particularly prone to sample bias, with points more often collected in areas highly accessible to humans (Bean et al. 2012), or in countries that upload their data to platforms like GBIF more consistently (Meyer et al. 2016). Bias can be controlled via a number of techniques, such as including covariates that act as a proxy for the bias (Warton et al. 2013), manipulating background points (Ranc et al. 2016, Vollering et al. 2019), thinning presence points across the geographic (Veloz 2009) or the environmental space (de Oliveira et al. 2014), or weighting data points (Elith et al. 2010). While sampling bias can be sometimes obvious when we compare our sample to the known approximate distribution of the species (e.g. by using IUCN range maps, or atlases), niche filling is harder to evaluate, as we only have good knowledge of the historical biogeography of a relatively small number of species. In many cases, the current distribution of species may result in a circular reasoning, where small ranges may suggest narrow climatic tolerance while the species only persists in a given geographic area for different reasons, e.g. because of anthropogenic impact (Di Marco and Santini 2016). Multiple studies have discussed the under-estimation of the niche due to historical range contractions (Varela et al., 2009; Maiorano et al., 2013; Martínez-Freiría et al., 2016; Faurby & Araújo, 2018), and demonstrated these may largely influence our future projections (Martínez-Freiría et al. 2016, Faurby and Araújo 2018). A possibility to alleviate this effect is using a multi-temporal approach (or time-calibrated models) by including historical data associated with the corresponding temporal climatic variables in the model training (Nogués-Bravo 2009, Maiorano et al. 2013). Yet, historical records are rarely available, so we should expect that the niche always tends to be under-estimated by an unknown extent compared to the true species potential, and climatic projections may therefore tend to be pessimistic on average about future species occurrence (Martínez-Freiría et al. 2016, Faurby and Araújo 2018).

### 4.7 Binarization

Accuracy metrics can fool us easily, and should not be used acritically to assess the reliability of a model, especially considering that they can provide higher estimates in sub-optimal conditions as we have shown here (Fig. S1). Some of the problems discussed above arise from the binarization of probabilistic model outputs into suitable and unsuitable areas (e.g. to determine the area of range contraction or expansion) based on a threshold. In fact, the true AUC tends to perform better than true TSS (Fig. S1-S2), and the estimated AUC has similar values to those of the true AUC under optimal conditions, whereas the estimated TSS is consistently higher than the true TSS, thus overestimating accuracy. Studies using MaxEnt typically only show AUC values (standard output of the software), even though model predictions are binarized. Here we show that while AUC is, as expected, highly correlated with TSS, high AUC can correspond to low TSS (Fig. S2). The problem arises from the fact that even when the model performs well, the threshold that maximizes discriminatory ability on the training dataset may not discriminate well true presence/absence, especially under different environmental conditions. Additionally, classical cross-validation is performed by using a split-sample approach, but a better and more informative option is to cross-validate on spatially independent samples (Bahn and McGill 2013, Roberts et al. 2017).

Additionally, other performance metrics focusing on probabilities (e.g. Boyce index) should be considered when possible. Several authors have argued that binarization should be entirely avoided unless it is clearly justified by the model application’s objective (Guillera-Arroita et al. 2015). Our results support this recommendation, and actually indicate that binary outputs should never be considered or used to quantify changes in distribution areas. Alternative approaches to summarize the results should be considered, such as looking at trends in predicted probabilities per areas.

### 4.8 Additional sources of uncertainty to be considered

In this study we evaluated the sensitivity of SDM predictions to a number of modelling settings and common violations of SDM assumptions. However, there are additional factors that we did not consider that can further contribute to making predictions less reliable. These include the spatial accuracy of data points in relation to the resolution used (Graham et al. 2008), the taxonomic accuracy of the data points (i.e., species confused with others, especially from citizen science data), and the ambiguous taxonomy of the species that may lead to merging data for different species, or viceversa not considering part of the distribution of a species (Araújo et al. 2019). Furthermore, species distribution models assume that the species niche is static, thereby ignoring intraspecific variation and local adaptations across populations (Pearman et al. 2010, Valladares et al. 2014).

This can be particularly problematic in climate change studies, as populations can adapt to climate change (Hoffmann and Sgró 2011), and different populations can hold diverse degrees of adaptation potential (Razgour et al. 2019).

### 4.9 Concluding remarks

Estimating the distribution of a species is a non-trivial task, as it requires a careful consideration of the biology of the species and its historical biogeography. Uncertainty is expected to be particularly high in studies modelling hundreds or thousands of species (Warren et al. 2013, 2018, Visconti et al. 2016, Newbold 2018, Thuiller et al. 2019), where species-specific considerations on the geographic extent or variables to include become impracticable, and normally the same geographic extent for sampling background points or set of variables is used. These studies are powerful for communicating important messages at the level of geographic areas (e.g., biomes) and entire communities, but need to be interpreted with extreme caution, and are ill-suited for drawing inferences at the level of species.

Our study indicates that our ability to predict future species distribution is low under on average, and can be low to the point of not being meaningful when conditions are far from optimal, especially when models’ predictions are binarized. Hence, SDM based climate change forecasting must adhere to the highest standards, must be clearly described (Zurell et al. 2020), and the estimated accuracy of models should be interpreted with extreme care, as well as the results, especially in relation to the quantification of range shifts, contraction and expansion, and the identification of areas that will be lost or gained. These considerations are also valid (and perhaps more problematic considering the wide temporal window and static niche assumption) in the case of hind-casting to paleoclimates, which is now common in studies focused on refugia and phylogeography (e.g. Svenning et al. 2011). Future research may focus on developing novel approaches to improve, synthesize and communicate SDM projections.

