## Supplementary materials for "Assessing the reliability of species distribution projections in climate change research"

`

### Appendix 1. Literature review on SDM modelling and projections

We conducted a literature review on common practices in species distribution modelling papers projecting models to a different time period. On the 10th of March 2020, we queried Web Of Science with the following string:

(("species distribut*" OR "habitat distribut*" OR "climat* envelope" OR "habitat suitab*" OR niche) AND (model*) AND ("climate change" OR "climate change scenario" OR future OR "range contraction" OR "range expansion" OR "range shift*") AND (Project* OR Predict*))

As similar reviews covering an earlier period exist (e.g. Fourcade et al. 2018), we limited the search to papers published between 2015 and 2019. This returned 4,157 hits, which, after removing duplicates records, included 3,927 hits.

Because the number of papers published in the topic has increased enormously over time and reviewing all of them was intractable and well beyond the scope of this paper, we sampled 50 papers per year and based our review on 250 papers (6.36% of the total). We scanned the 250 papers to assess if they were relevant, i.e. if they were research papers fitting correlative species distribution models and projecting to different times (or environmental space), thus including forecasting and hindcasting approaches. We excluded reviews, methodological papers and/or opinion papers. In total, we excluded 158 papers and retained 92 papers meeting our criteria (Table S1).

<u>For each of these papers we recorded:</u>
1) The sample size used for the modelling. In case of papers including multiple species we recorded the minimum sample size among all species used to fit the SDMs. If spatial or environmental thinning of data points was applied, we recorded the final sample size effectively used for the modeling.
2) If the entire set of 19 bioclimatic variables was used (or any other climatic set of variables) or specific variables were chosen.
6) If the selection of variables was explicitly justified, either on the basis of the species biology or on previous studies.
7) If the number of environmental variables was reduced using an approach based on variance inflation factors, correlation coefficients, or best fit to data, without additional biological considerations.
8) If a single model or an ensemble forecasting approach was used.
9) Which model(s) was/were employed.
10) If the continuous model outputs were binarized into presence/absence (suitable/unsuitable) either maximizing discrimination measures or using arbitrary thresholds.
11) If true absences were included, or pseudo-absences, background points, or presence-only models were used (e.g. BIOCLIM or DOMAIN).
12) What sampling approach for pseudo-absences or background points was used. Categories were: Bias = sampling that mimics sampling bias; Buffer = random sampling within a buffer around presence points; Distance-weighted = Sampling with higher intensity near (-) or far from (+) presence points; Globally = Random sampling globally; Outside climate envelope = Beyond climatic conditions observed for presence points; Study area = random sampling within a pre-defined study area.

We estimated the total number of relevant papers published in the 2015-2019 period using a bootstrapping approach. We sampled 250 values from binomial distribution with a proportion 0.364 of papers being relevant (92/250). We then multiplied the distribution of proportions for the total number of values and estimated the median and 95% interval of the distribution. This resulted in a median of 1429 (95CI 1194-1665) relevant articles published between 2015 and 2019.

`

### Supplementary figures

**Fig. S1.** AUC (a, b) and TSS (c, d) of the models fitted under optimal (n presences >200; niche completely filled; no environmental bias; no false predictors) and non-optimal conditions (n presences <50; 30% of niche filling; presences biased to 30% of the environmental gradient; no true predictor). Estimated = Estimated through internal cross-validation; Present and Future = True value validated against virtual reality for present and future; Contraction and Expansion = True value validated against virtual reality for predicted contraction and expansion areas; Dashed line = null expectation (no better than random); Dotted line = Value typically considered as "good" performance thresholds. The box edges are the 25th and 75th percentiles of the distribution, and whiskers 1.5 the inter-quartile range.

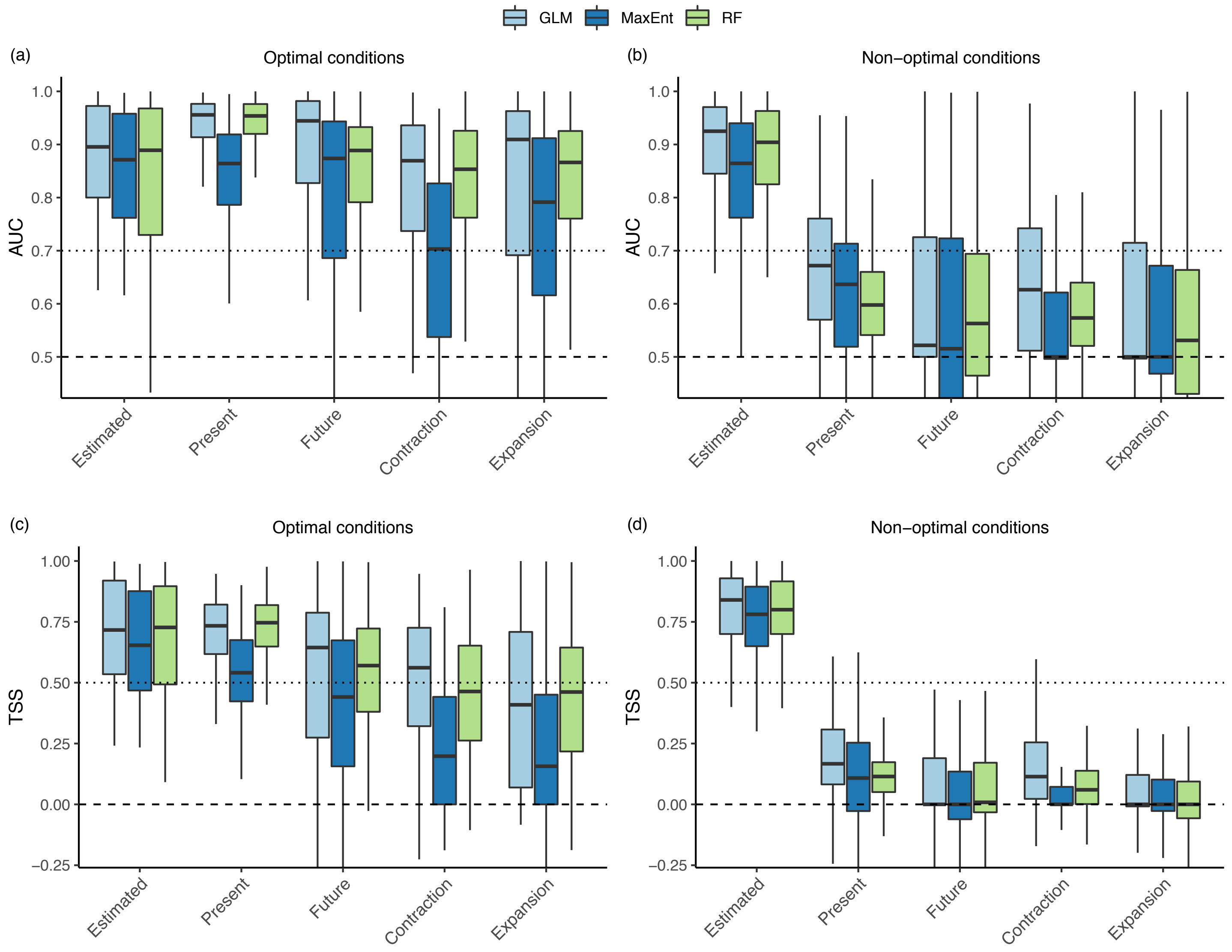

`

**Fig. S2.** Relationship between estimated (cv) and true (pres, fut) TSSs and AUCs for the three models.

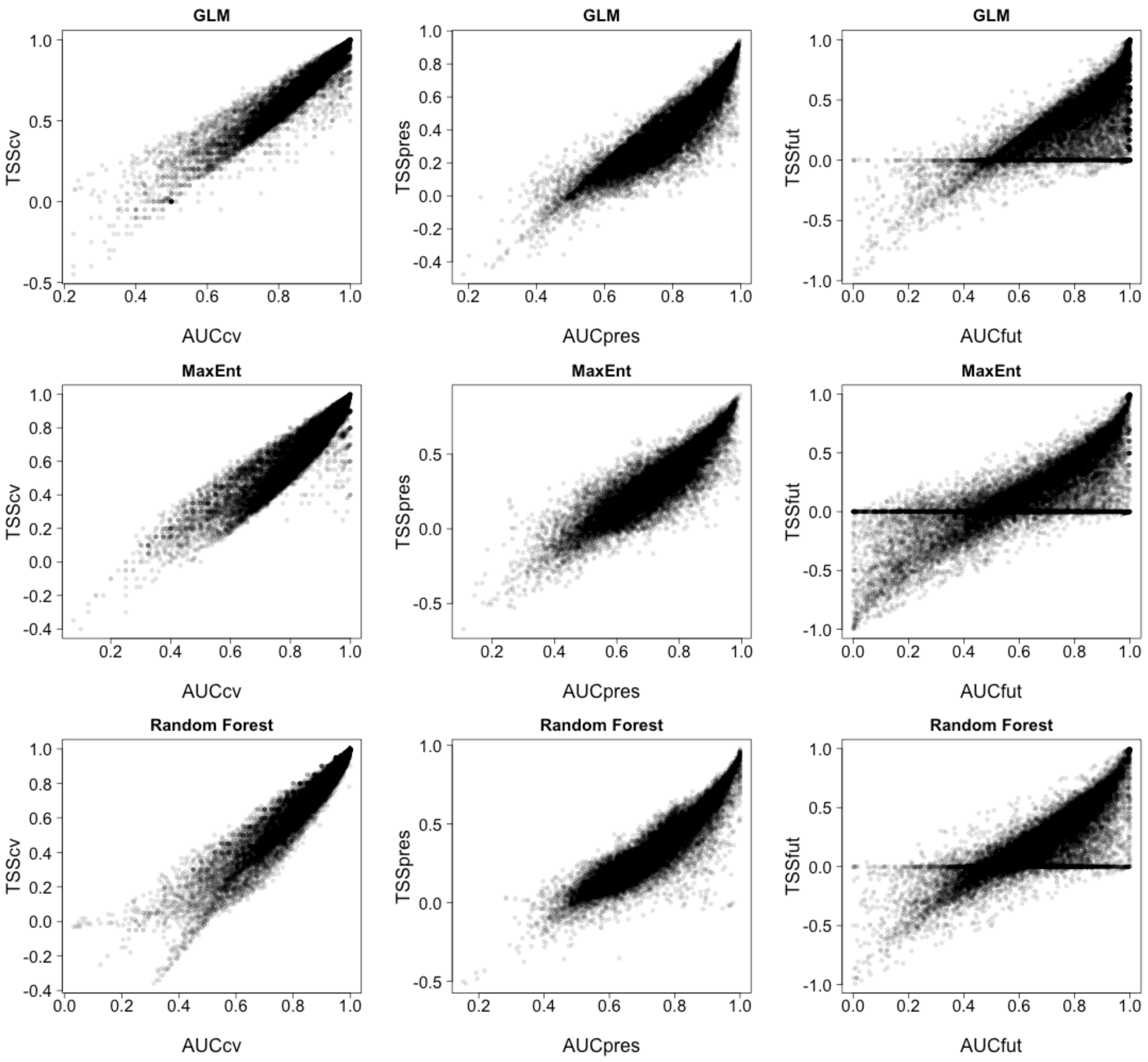

`

**Fig. S3.** Partial responses of the Random Forest model relating the estimated TSS by cross-validation ($TSS_{cv}$, green) and the true TSS for the present ($TSS_{pres}$, purple) and future ($TSS_{fut}$ orange) predictions with the treatments considered in the GLM. The partial response line represents the mean over 50 virtual species, whereas the shading the standard error around the mean.

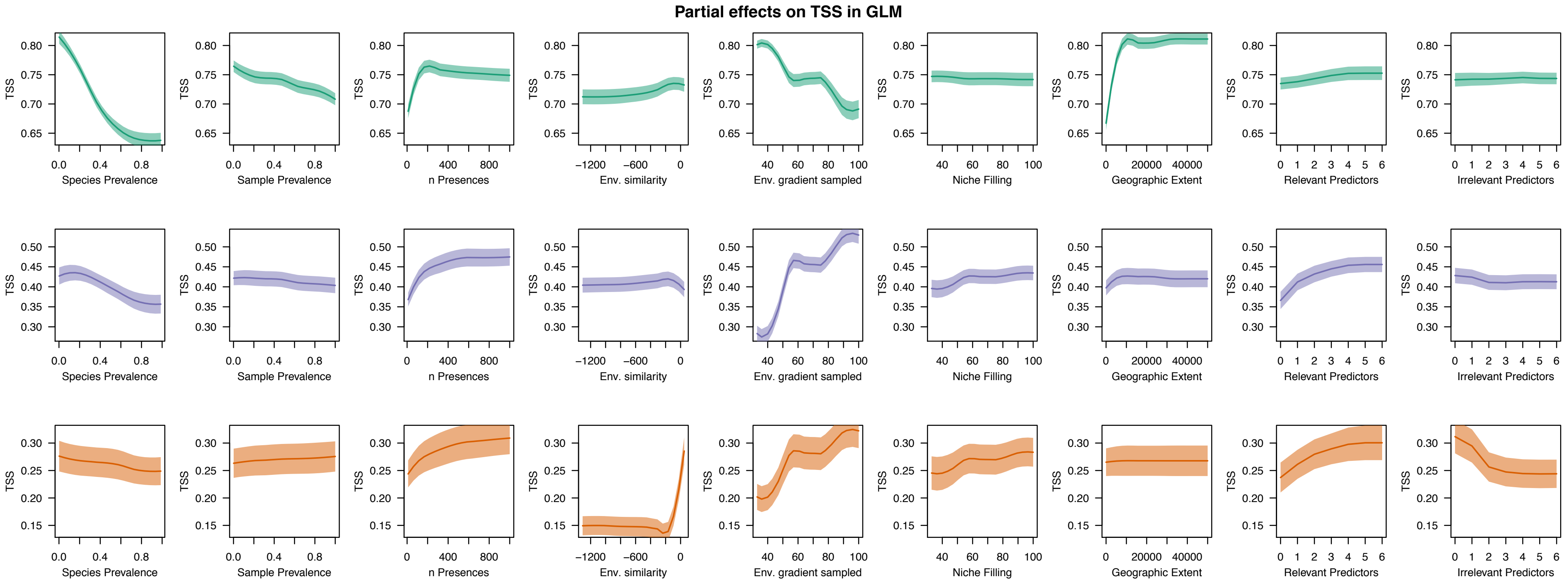

`

**Fig. S4.** Partial responses of the Random Forest model relating the estimated AUC by cross-validation ($AUC_{cv}$, green) and the true AUC for the present ($AUC_{pres}$, violet) and future ($AUC_{fut}$, orange) predictions with the treatments considered in the GLM. The partial response line represents the mean over all virtual species, whereas the shading the standard error around the mean.

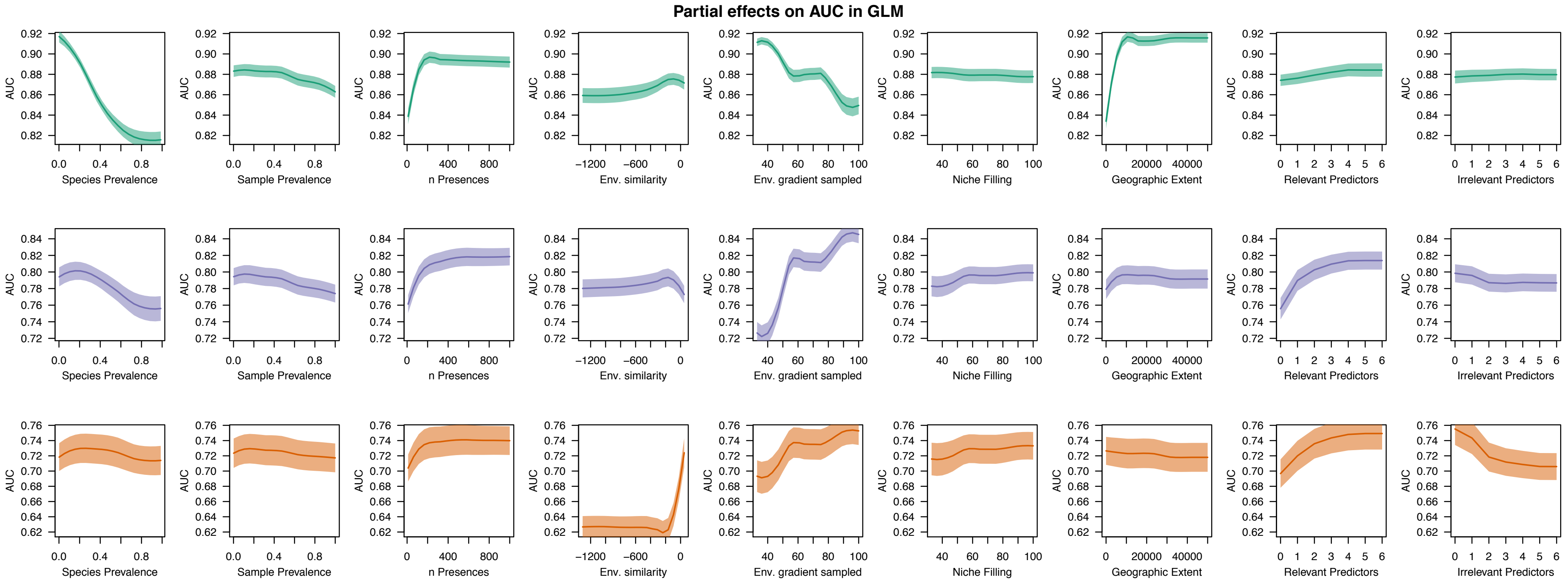

`

**Fig. S5.** Partial responses of the Random Forest model relating the estimated TSS by cross-validation ($TSS_{cv}$, green) and the true TSS for the present ($TSS_{pres,}$ purple) and future ($TSS_{fut}$ orange) predictions with the treatments considered in MaxEnt. The partial response line represents the mean over 50 virtual species, whereas the shading the standard error around the mean.

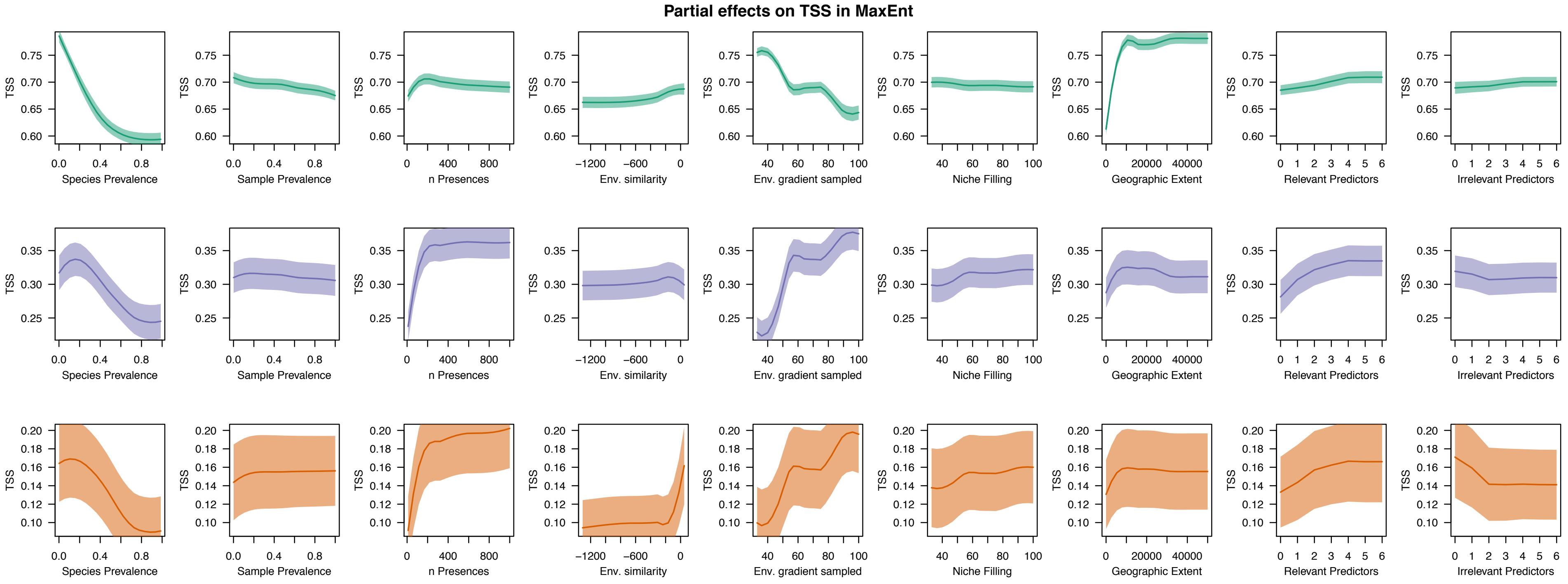

`

**Fig. S6.** Partial responses of the Random Forest model relating the estimated AUC by cross-validation ($AUC_{cv}$, green) and the true AUC for the present ($AUC_{pres}$, violet) and future ($AUC_{fut}$, orange) predictions with the treatments considered in MaxEnt. The partial response line represents the mean over all virtual species, whereas the shading the standard error around the mean.

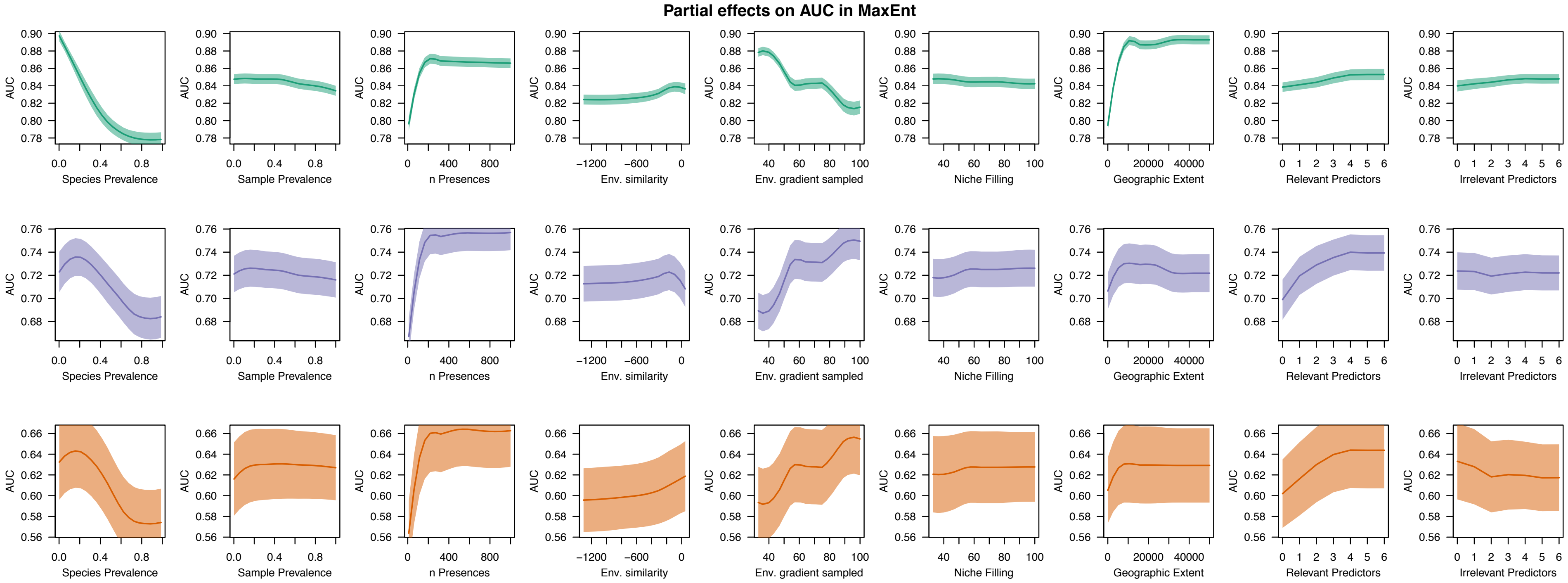

`

**Fig. S7.** Partial responses of the Random Forest model relating the estimated TSS by cross-validation ($TSS_{cv}$, green) and the true TSS for the present ($TSS_{pres}$, purple) and future ($TSS_{fut}$ orange) predictions with the treatments considered in the Random Forest. The partial response line represents the mean over 50 virtual species, whereas the shading the standard error around the mean.

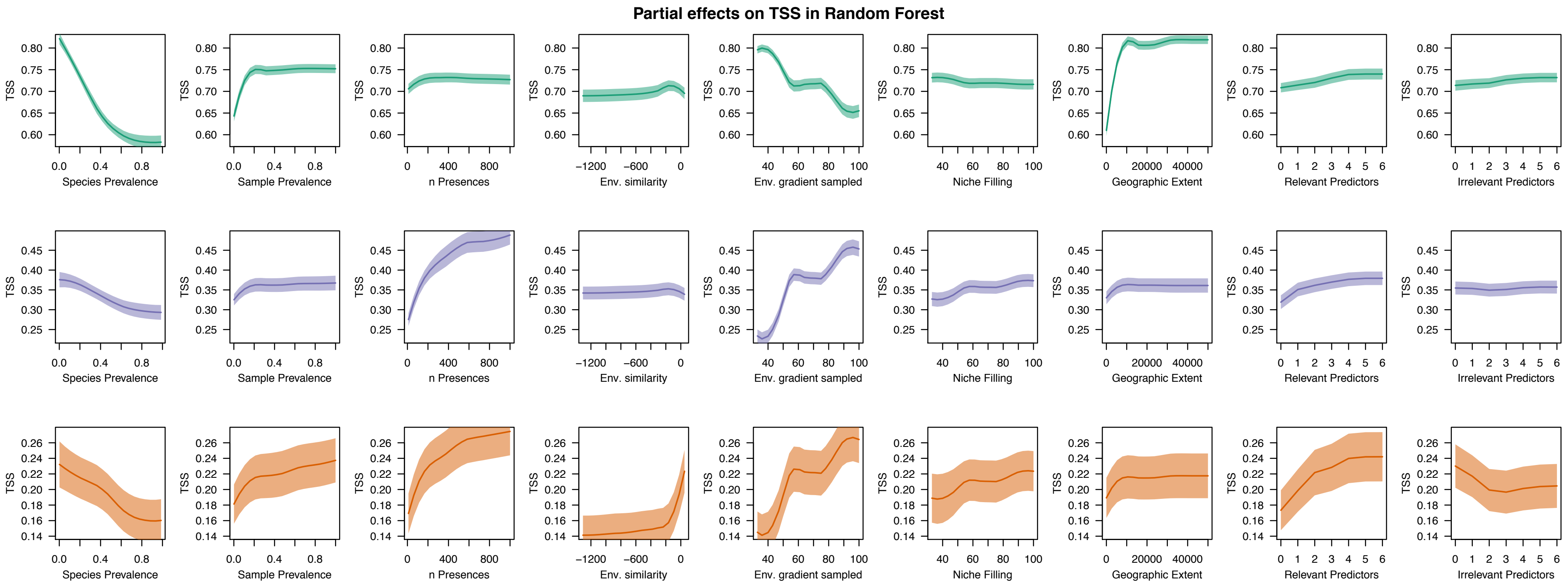

`

**Fig. S8.** Partial responses of the Random Forest model relating the estimated AUC by cross-validation ($AUC_{cv}$, green) and the true AUC for the present ($AUC_{pres}$, violet) and future ($AUC_{fut}$, orange) predictions with the treatments considered in the Random Forest. The partial response line represents the mean over 50 virtual species, whereas the shading the standard error around the mean.

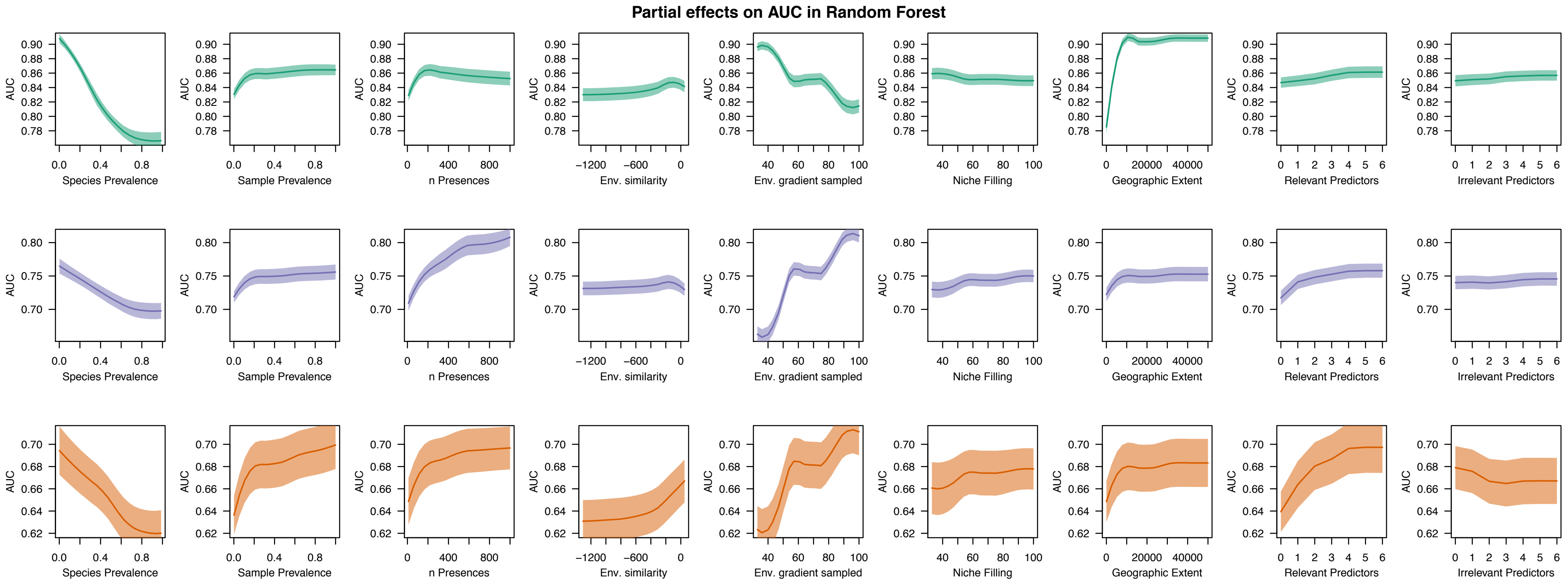

`

**Fig. S9.** Relative variable importance of different settings and conditions on the estimates of range expansion (a) and contraction (b) using binary outputs. Relative importance values are rescaled to 100 for each species. Bars represent the mean over all virtual species and error bars the standard error around the mean.

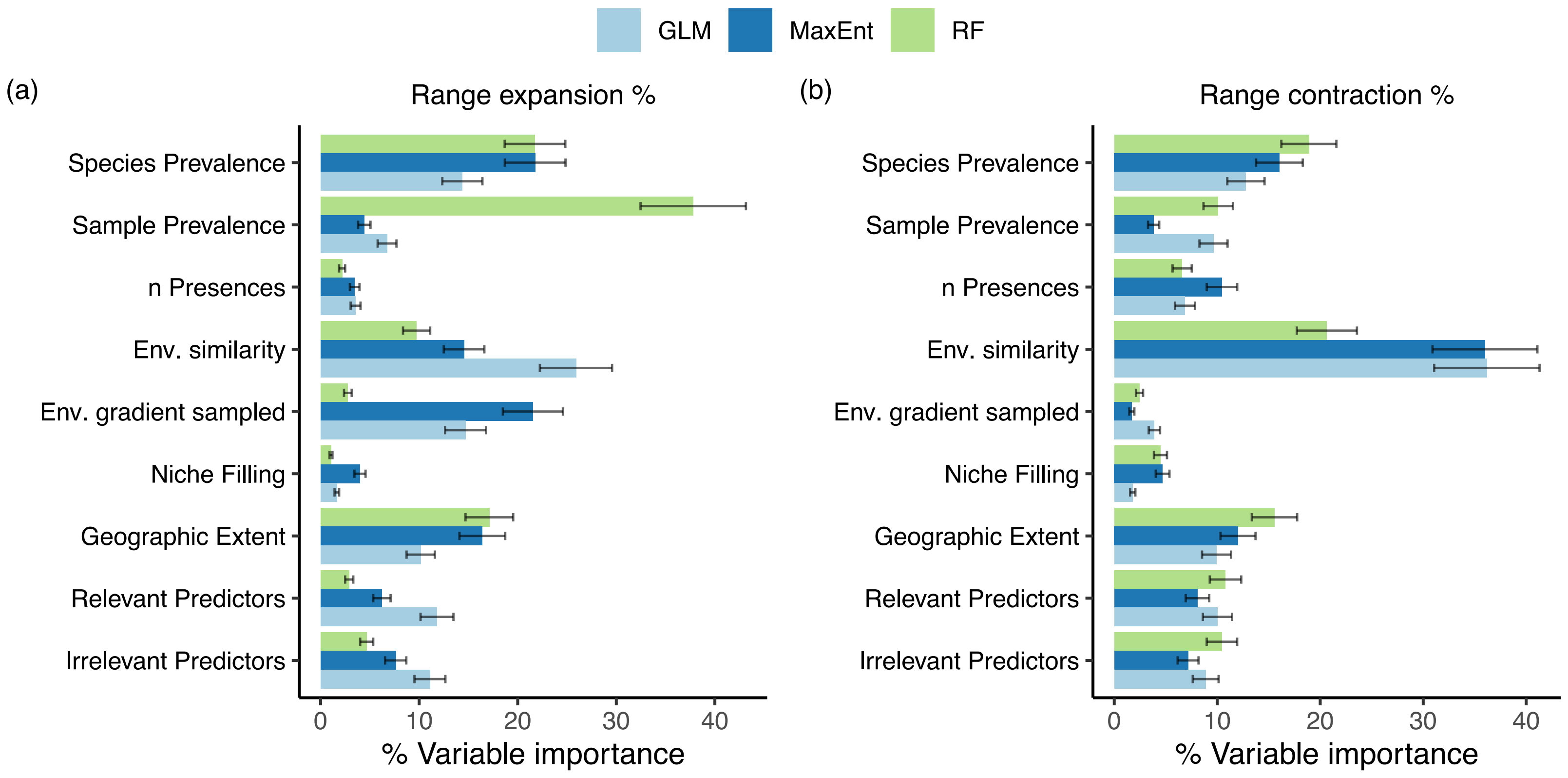

`

**Fig. S10.** Partial responses of the Random Forest model relating the estimated range contraction and expansion with the treatments considered in the GLM. The partial response line represents the mean over all virtual species, whereas the shading the standard error around the mean.

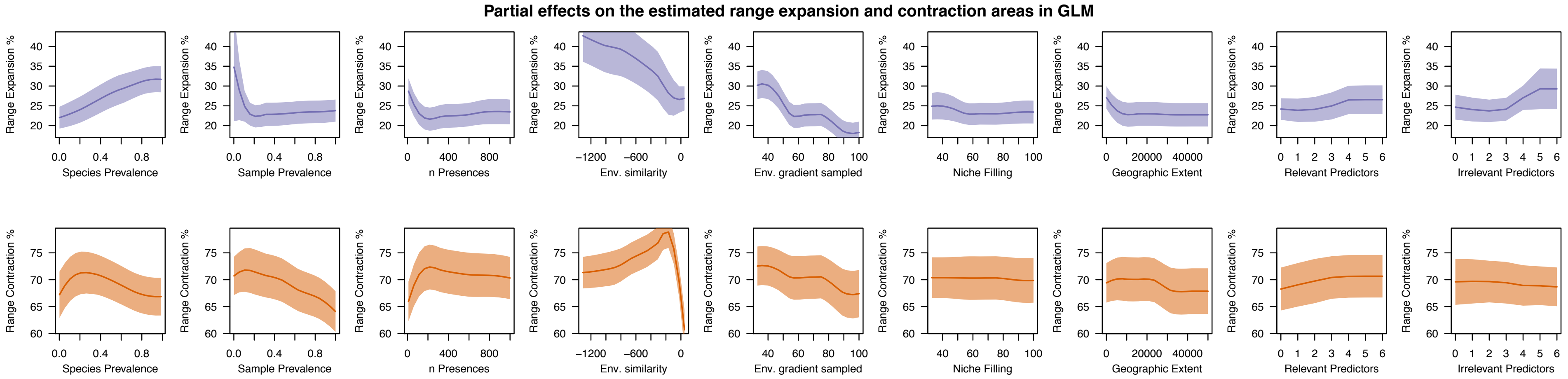

`

**Fig. S11.** Partial responses of the Random Forest model relating the estimated range contraction and expansion with the treatments considered in MaxEnt. The partial response line represents the mean over all virtual species, whereas the shading the standard error around the mean.

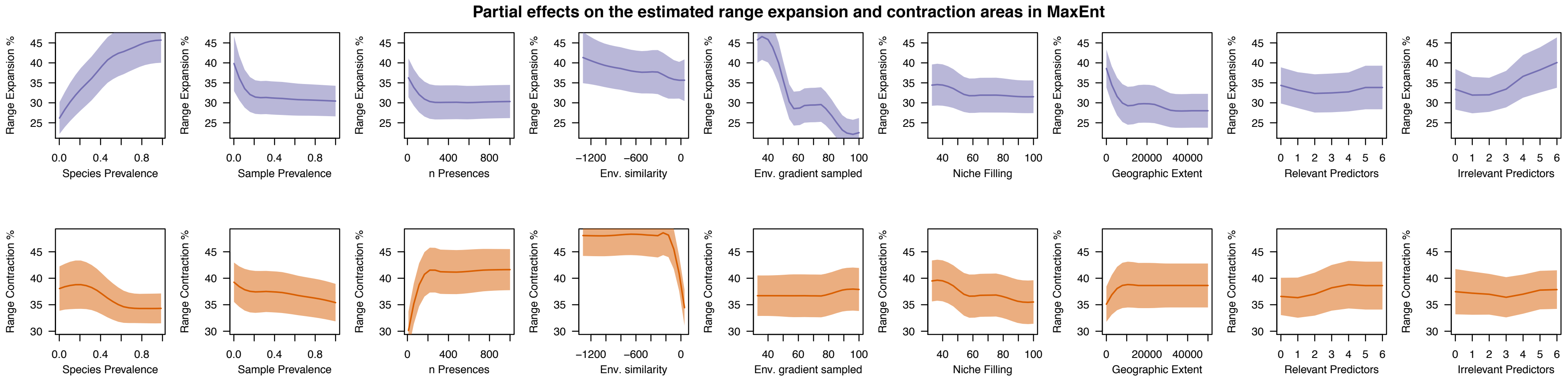

`

**Fig. S12.** Partial responses of the Random Forest model relating the estimated range contraction and expansion with the treatments considered in the Random Forest. The partial response line represents the mean over all virtual species, whereas the shading the standard error around the mean.

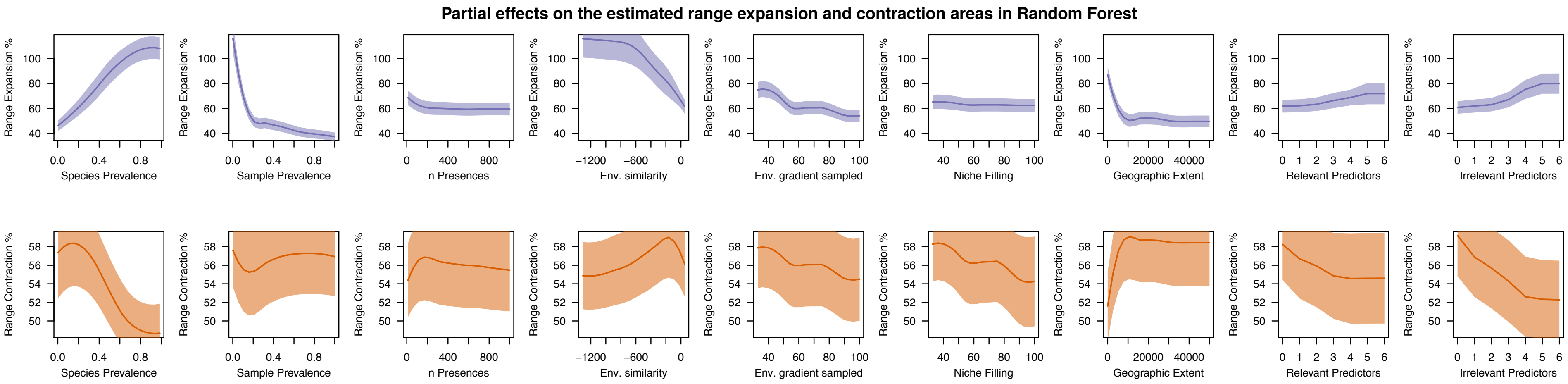
